# BrainEXP-NPD: a database of transcriptomic profiles of human brains of six neuropsychiatric disorders

**DOI:** 10.1101/2021.05.30.446363

**Authors:** Cuihua Xia, Teng Ma, Chuan Jiao, Chao Chen, Chunyu Liu

## Abstract

**Background:** Spatio-temporal gene expression has been widely used to study gene functions and biological mechanisms in diseases. Numerous microarray and RNA sequencing data focusing on brain transcriptomes in neuropsychiatric disorders have accumulated. However, their consistency, reproducibility has not been properly evaluated. Except for a few psychiatric disorders, like schizophrenia, bipolar disorder and autism, most have not been compared to each other for cross-disorder comparisons.

**Methods:** We organized 48 human brain transcriptome datasets from six sources. The original brain donors include patients with schizophrenia (SCZ, N=427), bipolar disorder (BD, N=312), major depressive disorder (MDD, N=219), autism spectrum disorder (ASD, N=53), Alzheimer’s disease (AD, N=765), Parkinson’s disease (PD, N=163) as well as controls as unaffected by such disorders (CTRL, N=6,378), making it a total of 8,317 samples. Raw data included multiple brain regions of both sexes, with ages ranging from embryonic to seniors. After standardization, quality control, filtering and removal of known and unknown covariates, we performed comprehensive meta- and mega-analyses, including gene differential expression and gene co-expression network.

**Results:** A total of 6922, 3011, 2703, 4389, 3507, 4279 significantly differentially expressed genes (FDR q < 0.05) were detected in the comparisons of 6 brain regions of SCZ-CTRL, 5 brain regions of BD-CTRL, 6 brain regions of MDD-CTRL, 4 brain regions of ASD-CTRL, 7 brain regions of AD-CTRL, and 6 brain regions of PD-CTRL, respectively. Most differentially expressed genes were brain region-specific and disease-specific. SCZ and BD have a maximal transcriptome similarity in striatum (ρ=0.42) among the four brain regions, as measured by Spearman’s correlation of differential expression log2 FC values. SCZ and MDD have a maximal transcriptome similarity in hippocampus (ρ=0.30) among the five brain regions. BD and MDD have a maximal transcriptome similarity in frontal cortex (ρ=0.45) among the five brain regions. Other disease pairs have a less transcriptome similarity (ρ<0.1) in all brain regions. PD is negatively correlated with SCZ, BD, and MDD in cerebellum and striatum. We also performed coexpression network analyses for different disorders and controls separately. We developed a database named BrainEXP-NPD (http://brainexpnpd.org:8088/BrainEXPNPD/), to provide a userfriendly web interface for accessing the data, and analytical results of meta- and mega-analyses, including gene differential expression and gene co-expression networks between cases and controls on different brain regions, sexes and age groups. Discussion: BrainEXP-NPD compiled the largest collection of brain transcriptomic data of major neuropsychiatric disorders and presented lists of differentially expressed genes and coexpression modules in multiple brain regions of six major disorders.

## Introduction

It is widely accepted that gene expression and gene-gene interaction are the basic tools to study and understand molecular normal and abnormal functions in organisms. Though genetic studies have revealed lots of disease risk variants, it still remains unclear how genetic factors impact on the disease. Transcriptome can help identify convergent molecular pathology^[1]^.

A number of public databases have been developed for studying expression variations and co-expression patterns in neuropsychiatric disorders, such as SZDB, AlzData, SMRIDB. SZDB is a specific database which focuses on SNP and gene annotation, spatio-temporal expression pattern analysis, network (PPI and co-expression) and pathway analysis, brain eQTL analysis and gene prioritization of schizophrenia^[2]^. AlzData covers high-throughput omic data, including genomics, transcriptomics, proteomics and functional genomics of Alzheimer’s disease^[3]^. However, both SZDB and AlzData only focus on a specific neuropsychiatric disease. SMRIDB (The Stanley Medical Research Institute Online Genomics Database) contains 988 samples from 12 studies and 6 platforms aimed at gene differential analysis and pathway/GO analysis of schizophrenia, bipolar, depression and healthy controls^[4]^. But cross-disorder analysis and co-expression analysis are not provided. Since these databases are constructed primarily on specific neuropsychiatric disorders or only gene expression, a more integrative data source and comprehensive research for neuropsychiatric disorders are urgently needed.

Here we developed a database which focuses on six complex neuropsychiatric disorders’ transcriptome profiles’ analyses. We first collected totally 8,317 brain samples from GEO (Gene Expression Omnibus), ArrayExpress and our own lab. It consists of six complex neuropsychiatric disorders (427 schizophrenia, 312 bipolar disorder, 163 Parkinson’s disease, 765 Alzheimer’s disease, 219 major depressive disorder, 53 autism spectrum disorder) and 6,378 controls (no neuropsychiatric-related disorders) across 55 brain regions from 48 individual datasets and different platforms (microarrays and RNA-seq). Then a consistent process workflow was applied to each individual dataset, and a combined-data analysis was conducted to uncover gene variations and gene co-expression patterns between cases and controls across different brain regions, sexes and age stages. Besides, a weighted gene coexpression network analysis (WGCNA)^[5]^ was carried out to figure out the disease-related modules and pathway enrichment. Finally, an integrative database of Transcriptomic Profiling in human Brains for six NeuroPsychiatric disorders (BrainEXP-NPD) was developed as a useful open free resource and reference for researchers and clinical workers.

## Material and Methods

### Study Design

The study was designed into three stages: Data Collection, Data Analysis and Visualization, as shown in Figure 1. We first collected the brain transcriptomic data from GEO^[6]^, ArrayExpress^[7]^, SMRIDB^[4]^, GTEx^[8]^, ROSMAP^[9]^ and PsychENCODE^[10]^. And then analyses were conducted for the collected data. Here we divided the analysis into two parts: differential gene expression (DEG) analysis, including individual dataset DEG analysis, DEG meta-analysis, DEG mega analysis; and gene co-expression analysis. We first processed each individual dataset according to the consistent workflow as the Figure 1 shows. And then, the datasets were sorted according to different brain regions. Next, each of the three types of DEG analyses was conducted across three aspects: brain region, sex and age stage. Besides, gene network analysis was conducted using weighted gene co-expression network analysis (WGCNA)^[5]^. In the third stage, we developed a database named Brain EXPression in NeuroPsychiatric Disorders (BrainEXP-NPD), which visualized the analysis results through various tables and figures.

**Figure 1.**
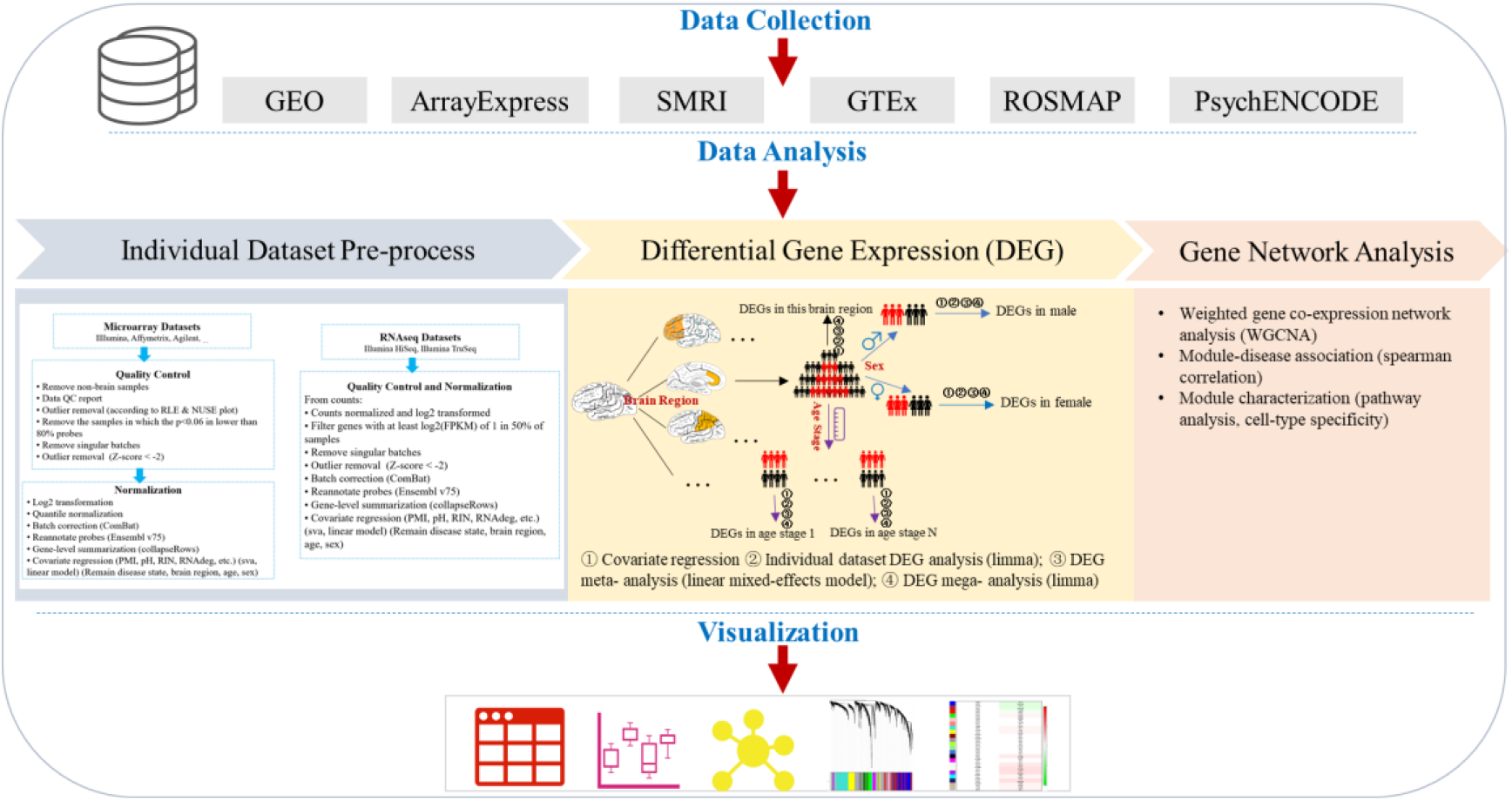
The study design of the BrainEXP-NPD database. DEGs: differential expressed genes

### Datasets

All the transcriptomic datasets included in the BrainEXP-NPD database are shown in Table 1. The data distributions across disease state, brain region, sex, age stage are shown in Figure 2. Totally 8,317 brain samples were collected from GEO, ArrayExpress and our own lab. It consists of six complex neuropsychiatric disorders (427 schizophrenia, 312 bipolar disorder, 163 Parkinson’s disease, 765 Alzheimer’s disease, 219 major depressive disorder, 53 autism spectrum disorder) and 6,378 controls (no neuropsychiatric-related disorders) across 55 brain regions from 48 individual datasets and different platforms (microarrays and RNA-seq) (Figure 2).

**Figure 2.**
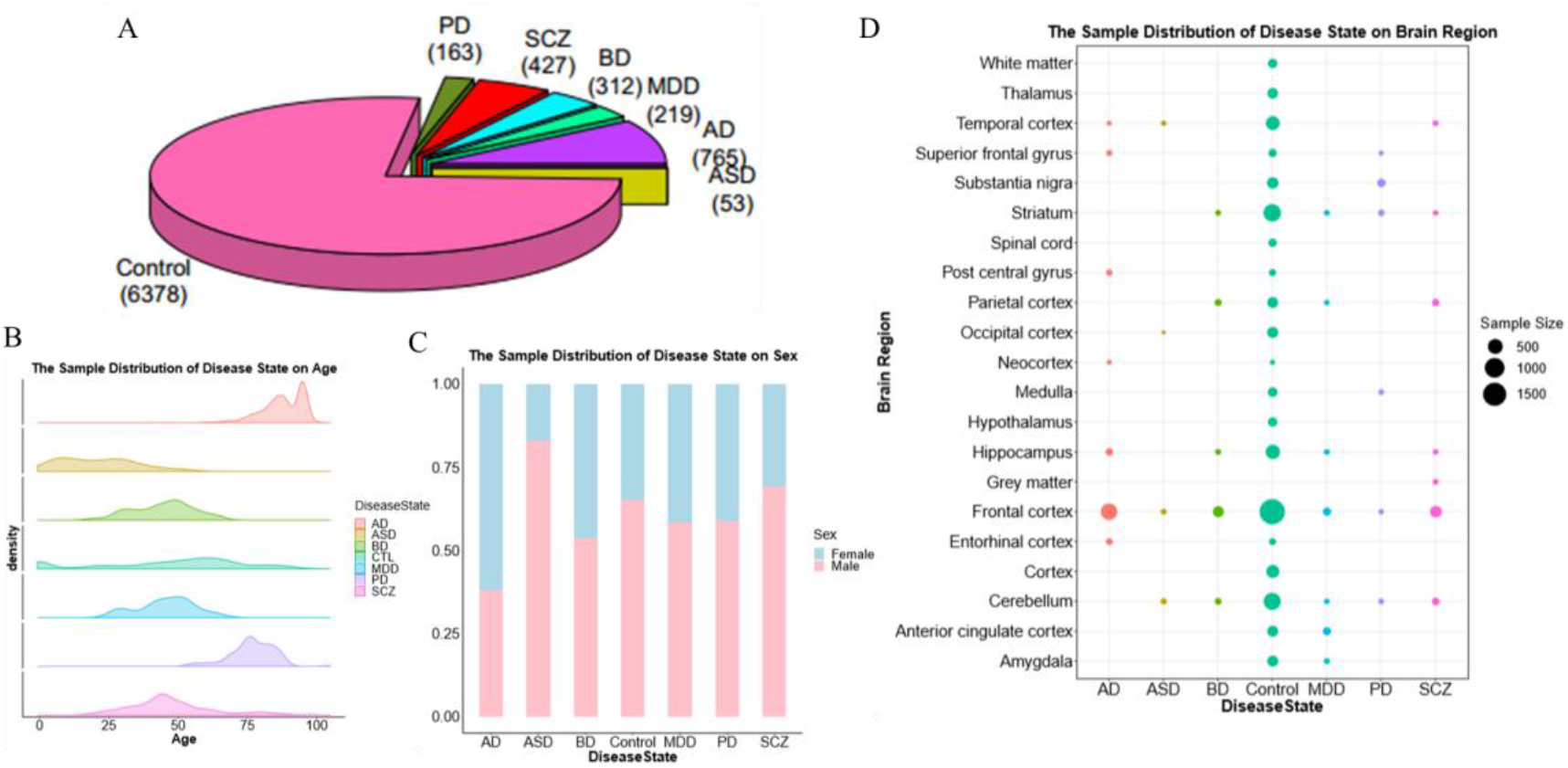
The data distribution in the BrainEXP-NPD database. A. The percentage of samples in different disease states. In BrainEXP-NPD, totally six neuropsychiatric disorders are included. B. The sample distribution in different ages of different disease states. C. The sample distribution in different sexes of different disease states. D. The sample distribution in different brain regions of different disease states.

**Table 1.** Number of individuals across disease state on transcriptome per brain project

| Name | # of Individuals |  | # of Brain regions | Disease state (# of individuals) |  |  |  |  |  |  |  |
| --- | --- | --- | --- | --- | --- | --- | --- | --- | --- | --- | --- |
|  | (# of samples) | Age |  | CTRL | SCZ | BD | MDD | ASD | AD | PD | Others |
| BrainCloud | 269 | Lifespan | 1 | 269 |  |  |  |  |  |  |  |
| BrainSpan | 42 | Lifespan | 6 | 42 |  |  |  |  |  |  |  |
| UKBEC | 134 | Postnatal | 6 | 134 |  |  |  |  |  |  |  |
| GTE<sub>x</sub> | 714 (2418) | Adult | 14 | 714 |  |  |  |  |  |  |  |
| BrainEXP | 2863 (4567) | Lifespan | 56 | 4,567 |  |  |  |  |  |  |  |
| CMC | 621 | Postnatal | 1 | 291 | 275 | 47 |  |  |  |  | 8 |
| BrainSeq | 746 | Lifespan | 3 | 341 | 184 | 69 | 152 |  |  |  |  |
| ROSMAP | 748 | Adult | 1 | 432 |  |  |  |  | 316 |  |  |
| SZDB2.0 | 537 | Postnatal | 1 | 279 | 258 |  |  |  |  |  |  |
| AlzData | 1246 |  | 4 | 562 |  |  |  |  | 684 |  |  |
| SMRIDB | 979 |  | 5 |  |  |  |  |  |  |  |  |
| PsychENCODE | 1695 | Postnatal | 2 | 899 | 531 | 217 |  | 48 |  |  |  |

**Table 1.** All the transcriptomic datasets included in the BrainEXP-NPD database

| Accession | Platform | Disease State | # of Disease Samples | # of CTL Samples | # of Genes |
| --- | --- | --- | --- | --- | --- |
| GSE4036 | HG-U133P | SCZ | SCZ 11 | 14 | 22601 |
| GSE7621 | HG-U133P | PD | PD 11 | 6 | 22601 |
| GSE17612 | HG-U133P | SCZ | SCZ 25 | 21 | 22601 |
| GSE21138 | HG-U133P | SCZ | SCZ 27 | 28 | 22601 |
| GSE21935 | HG-U133P | SCZ | SCZ 21 | 17 | 22601 |
| GSE48350 | HG-U133P | AD | AD 75 | 166 | 22601 |
|  |  |  | BD 51, |  |  |
| GSE53987 | HG-U133P | BD, MDD, SCZ | MDD 49, | 53 | 22601 |
|  |  |  | SCZ 43 |  |  |
| GSE54565 | HG-U133P | MDD | MDD 15 | 16 | 22601 |
| GSE54567 | HG-U133P | MDD | MDD 14 | 13 | 22601 |
| GSE54568 | HG-U133P | MDD | MDD 13 | 13 | 22601 |
| GSE54571 | HG-U133P | MDD | MDD 10 | 11 | 22601 |
| GSE54572 | HG-U133P | MDD | MDD 12 | 9 | 22601 |
|  |  |  | BD 21, SCZ |  |  |
| study5 | HG-U133P | BD, SCZ | 18 | 18 | 22601 |
| GSE5392 | HG-U133A | BD | BD 32 | 35 | 13252 |
| GSE8397-GPL96 | HG-U133A | PD | PD 28 | 17 | 13252 |
| GSE8397-GPL97 | HG-U133B | PD | PD 28 | 17 | 10413 |
| GSE20291 | HG-U133A | PD | PD 15 | 17 | 13252 |
| GSE20164 | HG-U133A | PD | PD 6 | 4 | 13252 |
| GSE20314 | HG-U133A | PD | PD 4 | 2 | 13252 |
| GSE54570 | HG-U133A | MDD | MDD 7 | 12 | 13252 |
| GSE54575 | HG-U133A | MDD | MDD 11 | 10 | 13252 |
|  |  |  | BD 11, |  |  |
| study2 | HG-U133A | BD, SCZ,<br>Depression | Depression<br>10, SCZ 18 | 27 | 13252 |
|  |  |  | BD 27, SCZ |  |  |
| study3 | HG-U133A | BD, SCZ | 27 | 28 | 13252 |
|  |  |  | BD 29, SCZ |  |  |
| study7 | HG-U133A | BD, SCZ | 35 | 32 | 13252 |
| GSE12685 | HG-U133A | AD | AD 5 | 5 | 13252 |
| GSE22570 | HG-1_0st | CTL | 0 | 48 | 23823 |
|  |  |  | BD 73, |  |  |
| GSE35978 | HG-1_0st | BD, SCZ,<br>Depression | Depression<br>25, SCZ 94 | 94 | 23823 |
| GSE36980 | HG-1_0st | AD | AD 25 | 44 | 23823 |
| GSE45878 | HG-1_1st | CTL | 0 | 196 | 23847 |
| GSE71620 | HG-1_1st | CTL | 0 | 385 | 23847 |
| GSE30453 | HuEx-1_0-st | CTL | 0 | 86 | 15927 |
| GSE30422 | HuEx-1_0-st | CTL | 0 | 86 | 15927 |
| GSE37263 | HuEx-1_0-st | AD | AD 7 | 8 | 15927 |
| GSE60862(UKBEC) | HuEx-1_0-st | CTL | 0 | 1163 | 15927 |
| GSE25219-GPL5175 | HuEx-1_0-st | CTL | 0 | 1268 | 15927 |
|  | Illumina |  |  |  |  |
|  | HumanRef-8 |  |  |  |  |
| GSE28521 | v3.0 expression<br>beadchip | ASD | ASD 36 | 38 | 17686 |
| GSE26927 | Illumina<br>HumanRef-8<br>v2.0 expression<br>beadchip | AD, PD, SCZ | AD 18, PD<br>19, SCZ 18 | 0 | 18190 |
| GSE28894 | Illumina<br>HumanRef-8<br>v2.0 expression<br>beadchip | PD | PD 52 | 57 | 15761 |
| GSE29378 | Illumina<br>HumanHT-12<br>V3.0 expression<br>beadchip | AD | AD 30 | 31 | 37804 |
| GSE37205(Illumina<br>Probe Annotation) | Illumina<br>HumanHT-12<br>V3.0 expression<br>beadchip | CTL | 0 | 96 | 25531 |
| GSE54562 | Illumina<br>HumanHT-12<br>V3.0 expression<br>beadchip | MDD | MDD 9 | 9 | 25531 |
| GSE54563 | Illumina<br>HumanHT-12<br>V3.0 expression<br>beadchip | MDD | MDD 25 | 24 | 25531 |
| GSE54564 | Illumina<br>HumanHT-12<br>V3.0 expression<br>beadchip | MDD | MDD 19 | 20 | 25531 |
| GSE38322(SubSeries of<br>GSE38609, methyl) | Illumina<br>HumanHT-12<br>V4.0 expression<br>beadchip | ASD | ASD 17 | 18 | 31426 |
| GSE30272(BrainCloud) | Illumina Human<br>49K Oligo array<br>(HEEBO-7 set) | CTL | 0 | 269 | 17161 |
| ROSMAP | non-stranded,<br>Illumina HiSeq | AD | AD 605 | 7 | 16263 |
| GTEx | non-stranded,<br>Illumina TruSeq<br>library<br>construction<br>protocol | CTL | 0 | 1629 | 19449 |
| BrainGVEX | non-stranded, | SCZ, BD | SCZ 90, BD<br>68 | 233 | 16660 |
|  | Illumina |  |  |  |  |
|  | HiSeq2000 |  |  |  |  |

### Identification of Differentially Expressed Genes

Limma package^[11]^ was used for the individual dataset DEG analysis and DEG mega analysis. For the meta-analysis, consistent differential expressed genes between cases and controls were figured out using linear mixed-effects model according to Michael J. Gandal’s workflow^[12]^. The significant threshold for DEGs was FDR q<0.05.

### Identification of Co-expressed Modules

Weighted gene co-expression network analysis (WGCNA)^[5]^ was used for identifying the co-expressed modules between cases and controls. Spearman correlation analysis was conducted between module eigengene and disease state to find disease related modules. The significant threshold for correlation was FDR q<0.05.

### Visualization of Analysis Results

The main languages for building the website are MySQL, HTML and Java. Apache was used for the server environment. Traditional MVC structure was used for the web design, the front-end page is implemented by the mainstream HTML+CSS+JS language combination, the Bootstrap component is used to customize the style of the page. Java language is used for the back-end to implement the logical functions of the website, and ECharts^[13]^ was used to display the data in a more intuitive way. The underlying layer uses MySQL to build and store our data.

## Results

### Differentially Expressed Genes

Totally 6922, 3011, 2703, 4389, 3507, 4279 significantly differentially expressed genes (FDR q < 0.05) were detected in the comparisons of 6 brain regions of SCZ-CTRL, 5 brain regions of BD-CTRL, 6 brain regions of MDD-CTRL, 4 brain regions of ASD-CTRL, 7 brain regions of AD-CTRL, and 6 brain regions of PD-CTRL, respectively. The numbers of significantly expressed genes of different brain regions between the six disorders and controls were shown in table 2.

**Table 2.**
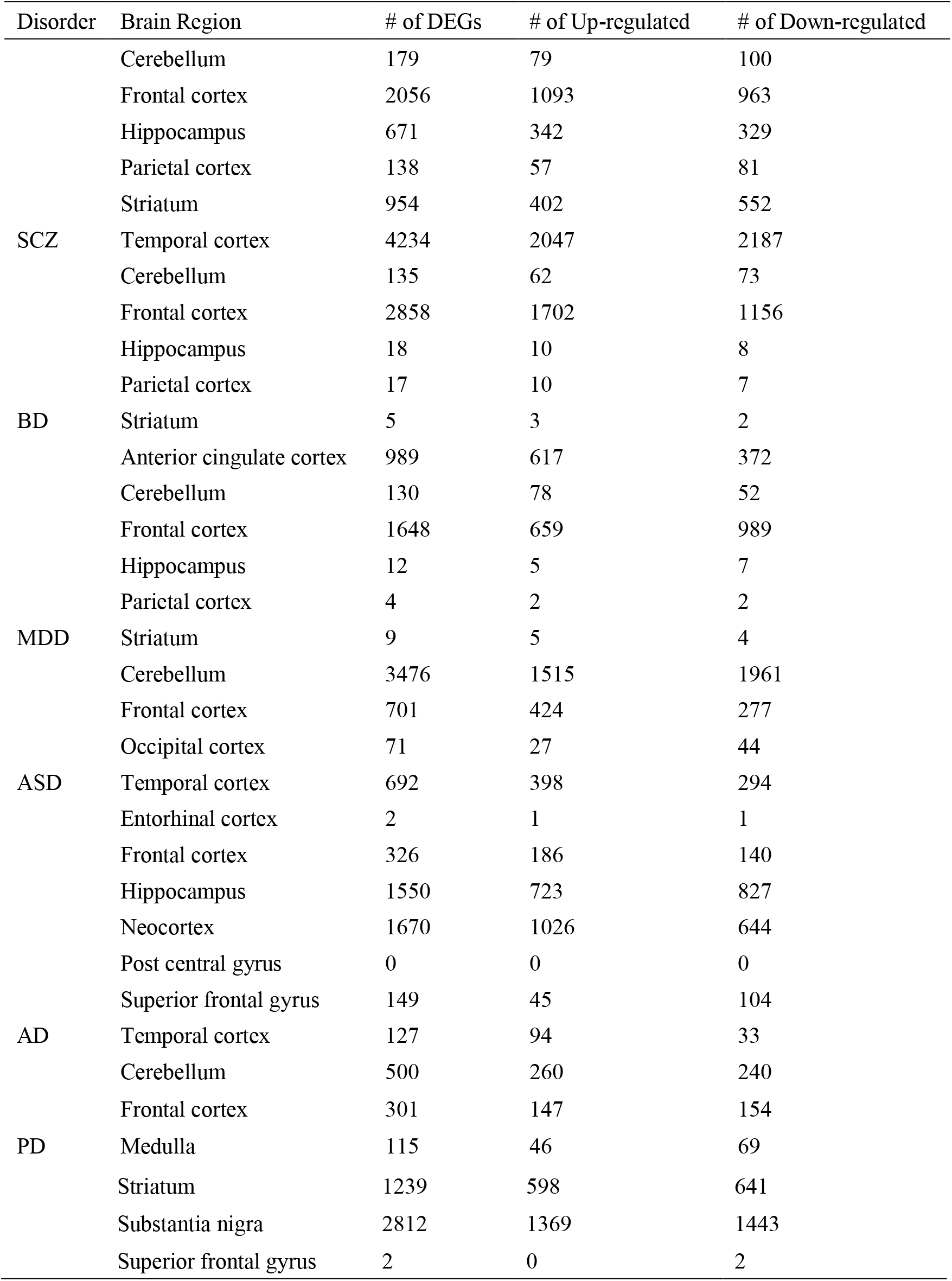
Numbers of significantly expressed genes of different brain regions between the six disorders and controls

**Table 3.**
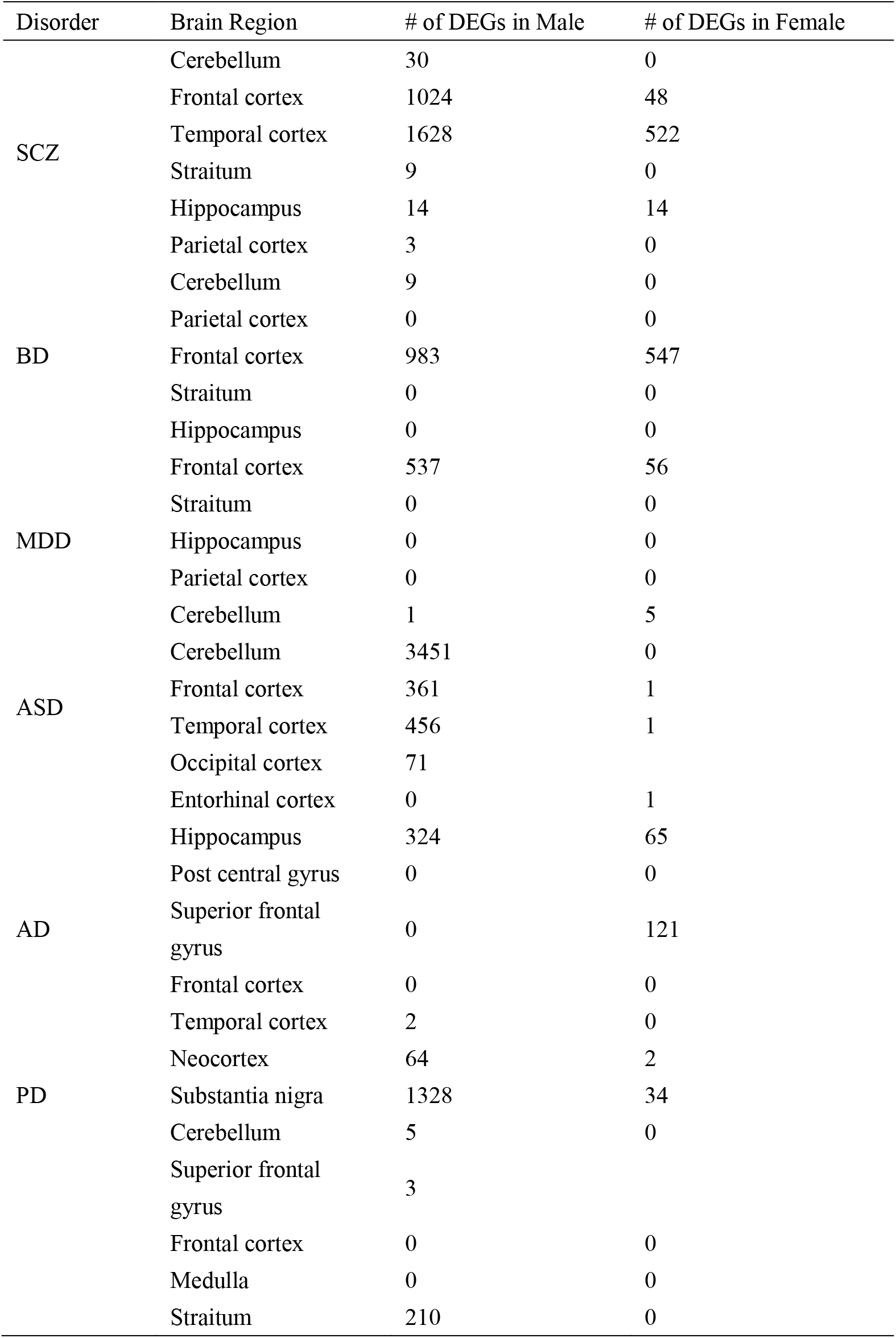
Numbers of significantly expressed genes of different sexes in different brain regions between the six disorders and controls

**Table 4.**
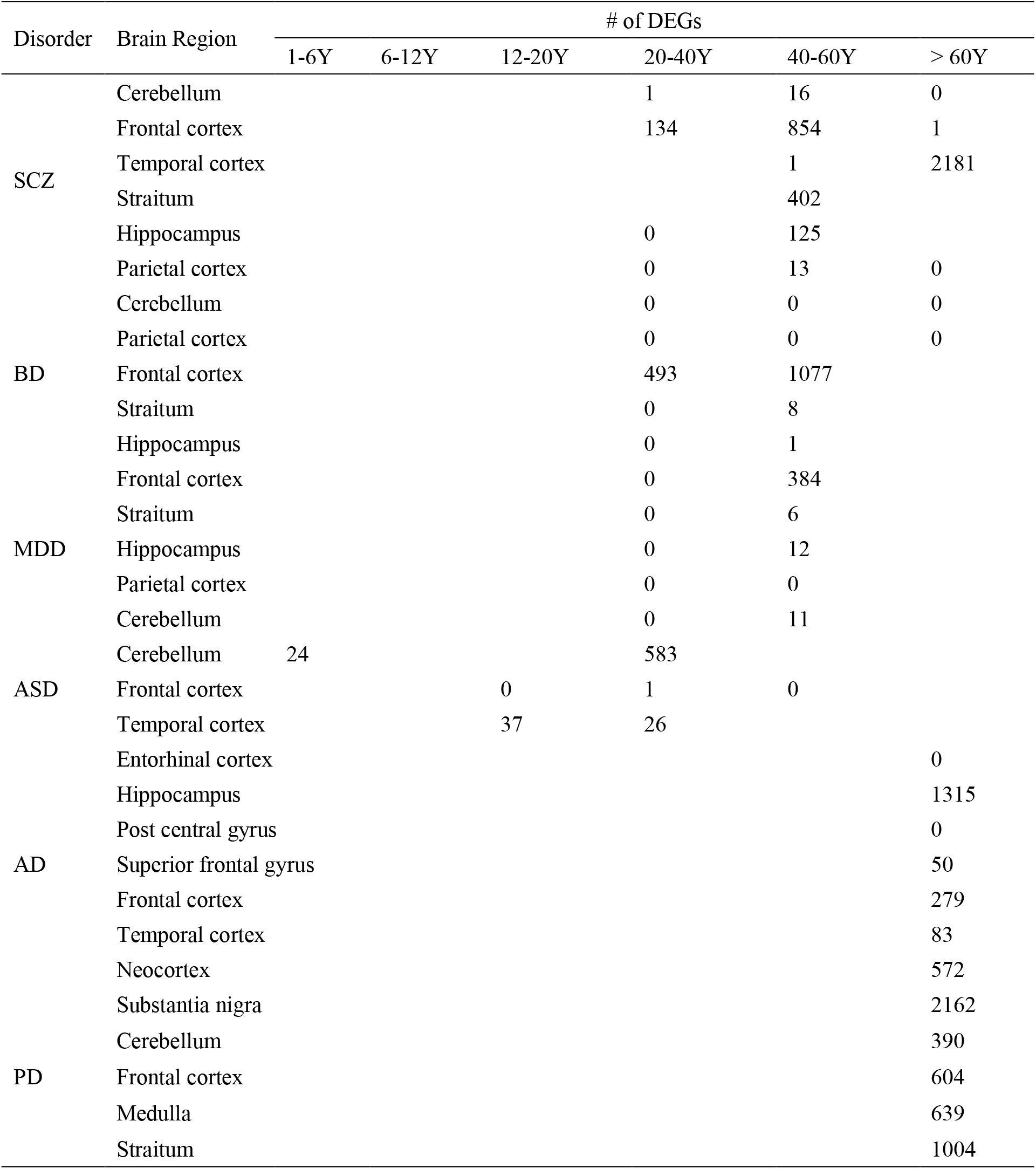
Numbers of significantly expressed genes of different age stages in different brain regions between the six disorders and controls

### Co-expressed Modules

Totally 4, 0, 0, 8, 13, 7 disease-related modules (FDR q < 0.05) were detected in SCZ-CTRL, BD-CTRL, MDD-CTRL, ASD-CTRL, AD-CTRL, and PD-CTRL dataset, respectively. The numbers of co-expressed modules detected between the six disorders and controls are shown in table 5.

**Table 5.**
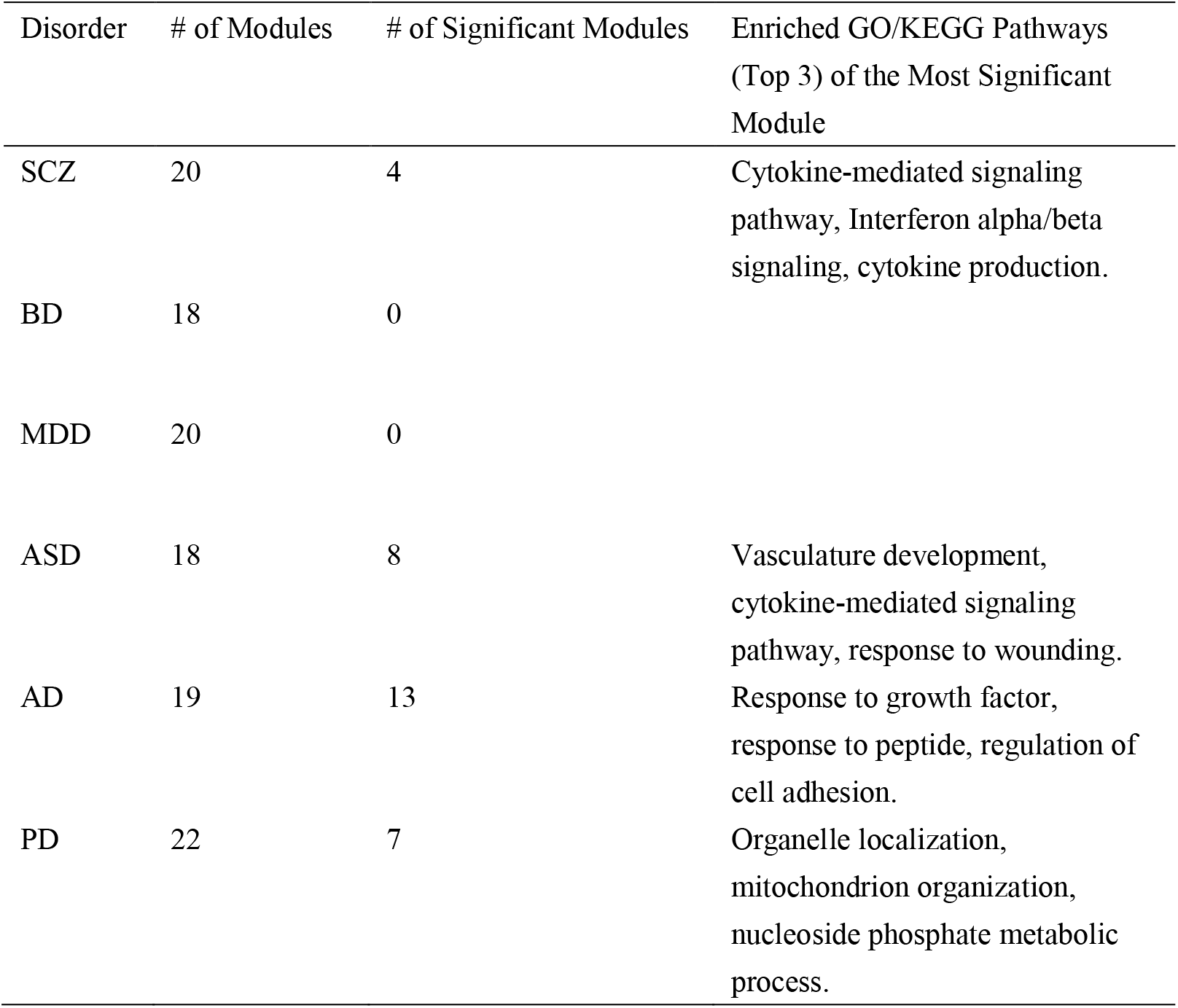
Numbers of co-expressed modules detected between the six disorders and controls

| Disorder | # of Modules | # of Significant Modules | Enriched GO/KEGG Pathways<br>(Top 3) of the Most Significant<br>Module |
| --- | --- | --- | --- |
| SCZ | 20 | 4 | Cytokine-mediated signaling pathway, Interferon alpha/beta signaling, cytokine production. |
| BD | 18 | 0 |  |
| MDD | 20 | 0 |  |
| ASD | 18 | 8 | Vasculature development, cytokine-mediated signaling pathway, response to wounding. |
| AD | 19 | 13 | Response to growth factor, response to peptide, regulation of cell adhesion. |
| PD | 22 | 7 | Organelle localization, mitochondrion organization, nucleoside phosphate metabolic process. |

### Visualization

The BrainEXP-NPD database (http://brainexpnpd.org:8088/BrainEXPNPD/index.html) was developed in an easy-to-use mode. We provide the DEG results and co-expression results on three aspects and six neuropsychiatric disorders. The result page for single gene search was shown in Figure 3. It includes totally four parts: Gene Basic Information, Gene Normalized Expression, Differential Gene Expression Summary Statistics, and Gene Co-expression. Here, we took human DRD2 as an example to demonstrate the usage of BrainEXP-NPD database, as shown in Figure 4.

**Figure 3.**
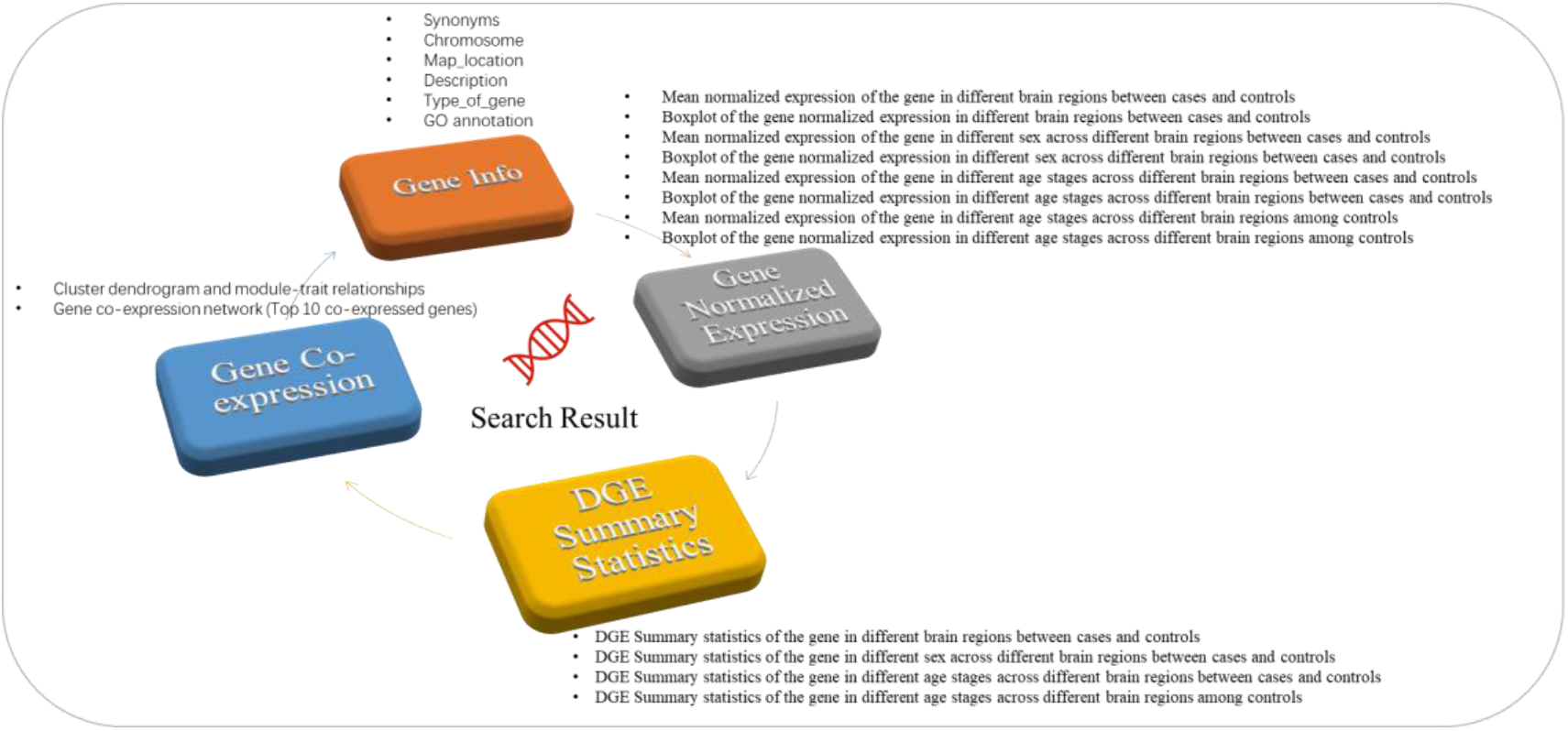
Data visualization content of the BrainEXP-NPD database.

**Figure 4.**
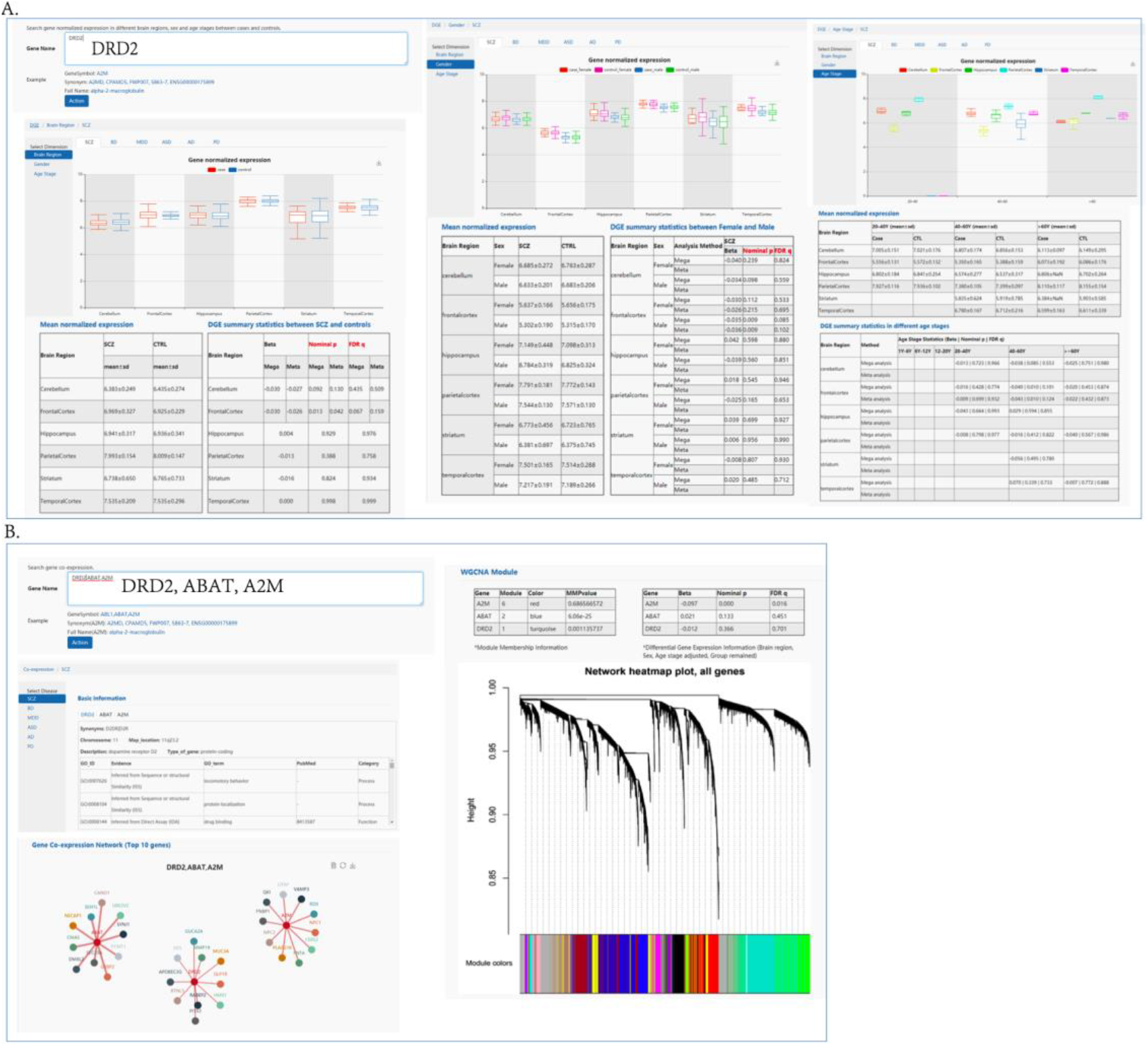
Search results and display. A. Gene differential expression retrieval results. B. Gene co-expression retrieval results.

## Summary and Perspectives

Currently, we provide a searchable & downloadable web entrance for results of normalized brain gene expression profiling, gene differential expression between cases and controls and co-expression. More microarray and RNA-seq data for brain expression of the major neuropsychiatric disorders are under continuously updating.

## Data Availability

The software used for data processing in this study is R 3.5.0, and the code for data analysis is updated on https://github.com/CuihuaXia/BrainEXP-NPD. All the data information can be freely accessed on http://brainexpnpd.org:8088/BrainEXPNPD/index.html.

## Authors’ contributions

Chunyu Liu and Chao Chen conceived, designed, and supervised the study. Cuihua Xia performed the overall analysis. Teng Ma developed the database webserver. Chuan Jiao contributed to the data collection and data analysis. Guihu Zhao contributed to develop the database webserver. Cuihua Xia wrote the manuscript. All authors read and approved the final manuscript.

## Acknowledgement

This work was supported by National Natural Science Foundation of China grants 81401114, 31571312, the National Key Plan for Scientific Research and Development of China (2016YFC1306000), Innovation-Driven Project of Central South University (No. 2015CXS034, 2018CX033) (to C. Chen), and NIH grants 1 U01 MH103340-01, 1R01ES024988 (to C. Liu). All the data contributors are sincerely appreciated for data submitted in the GEO and other databases. We are grateful to Professor Jinchen Li and National Clinical Research Center for Geriatric Disorders, Xiangya Hospital, Central South University for hosting our webserver.

